# Loss of Trem2 in microglia leads to widespread disruption of cell co-expression networks in mouse brain

**DOI:** 10.1101/248757

**Authors:** Guillermo Carbajosa, Karim Malki, Nathan Lawless, Hong Wang, John W. Ryder, Eva Wozniak, Kristie Wood, Charles A. Mein, Richard J.B. Dobson, David A. Collier, Michael J. O’Neill, Angela K. Hodges, Stephen J. Newhouse

## Abstract

Rare heterozygous coding variants in the Triggering Receptor Expressed in Myeloid cells 2 (TREM2) gene, conferring increased risk of developing late-onset Alzheimer's disease, have been identified. We examined the transcriptional consequences of the loss of Trem2 in mouse brain to better understand its role in disease using differential expression and coexpression network analysis of Trem2 knockout and wild-type mice. We generated RNA-Seq data from cortex and hippocampus sampled at 4 and 8 months. Using brain cell type markers and ontology enrichment, we found subnetworks with cell type and/or functional identity. We primarily discovered changes in an endothelial-gene enriched subnetwork at 4 months, including a shift towards a more central role for the Amyloid Precursor Protein (App) gene, coupled with widespread disruption of other cell-type subnetworks, including a subnetwork with neuronal identity. We reveal an unexpected potential role of Trem2 in the homeostasis of endothelial cells that goes beyond its known functions as a microglial receptor and signalling hub, suggesting an underlying link between immune response and vascular disease in dementia.

## 1 Introduction

Genome-wide association studies (GWAS) and genome sequencing have identified more than 25 Alzheimer’s disease (AD) risk loci, including common, low-risk variants and rare moderate risk variants, in addition to the classical risk variants in the APOE gene (R. Guerreiro et al., 2013; Hollingworth et al., 2011; Jonsson et al., 2013; Lambert et al., 2013; Naj et al., 2011; Pimenova et al., 2017; Steinberg et al., 2015). Although the identity of many of the associated disease genes and the mechanisms by which they increase risk remain unclear, there is evidence that they cluster around the immune system, protein and lipid metabolism, especially inflammatory response, endocytosis and APP metabolism (Hardy et al., 2014; Naj et al., 2017; Villegas-Llerena et al., 2016). Many implicated genes encode proteins which are highly expressed in microglia, (ABI3, PLCG3, TREM2, SPI1, BIN1, CD33, INPP5D, MS4A6A) and/or have a role in the innate immune system in the brain (Huang et al., 2017; Pimenova et al., 2017). One of these, TREM2 (Triggering Receptor Expressed On Myeloid Cells 2), has an AD associated risk allele (R47H) with an effect size in AD similar to that of ApoE4 (OR 2.90-5.05) (R. Guerreiro et al., 2013; Jonsson et al., 2013).

TREM2 function can be compromised as a result of rare non-synonymous variants which cause Nasu-Hakola disease when both alleles are affected (Numasawa et al., 2011; Ridha et al., 2004; Soragna, 2003) or significantly increase the risk of developing Alzheimer’s disease (AD) (R. Guerreiro et al., 2013; Jonsson et al., 2013; Sims et al., 2017), behavioral variant frontotemporal dementia (R. J. Guerreiro et al., 2013; Le Ber et al., 2013), semantic variant of primary progressive aphasia or occasionally Parkinson’s disease (PD) (Borroni et al., 2014; Liu et al., 2016; Rayaprolu et al., 2013) when one allele is affected. Evidence suggests TREM2 may be important for normal brain remodeling which peaks around adolescence (Chertoff et al., 2013) as well as in later life in response to age-related damage or pathologies. It isn’t clear whether the same signals or brain regions are affected by TREM2 activity throughout life. In people where TREM2 is compromised by complete loss of function, symptoms begin in adolescence and tend to implicate frontal lobe dysfunction, whereas in people with preservation of some normal TREM2 activity, symptoms appear much later in life and implicate the hippocampus (Giraldo et al., 2013). Although in both cases, progressive white matter changes in the brain and dementia symptoms occur.

TREM2 is a receptor highly expressed on macrophages, including microglia in the brain (Butovsky et al., 2014; Hickman et al., 2013; Paloneva et al., 2002; Schmid et al., 2002). It appears to be an important damage sensing receptor. It can respond to lipid and lipoprotein species such as phosphatidyl serine (Cannon et al., 2012; Wang et al., 2015), clusterin and apolipoproteins including APOE (Atagi et al., 2015; Poliani et al., 2015; Yeh et al., 2016) and nucleotides and anionic species such as heparin sulphate, proteoglycans or other negatively charged carbohydrates (Daws et al., 2003; Kawabori et al., 2015; Kober et al., 2016) and is required for efficient bacterial clearance (N’Diaye et al., 2009). TREM2 signalling is propagated through the adaptor protein DAP12 which activates a number of pathways including Syk, P13K and MAPK which culminate in increased phagocytosis and expression of an anti-inflammatory phenotype in microglia (Kleinberger et al., 2014; Neumann and Takahashi, 2007; Peng et al., 2010; Poliani et al., 2015; Takahashi et al., 2007). Loss of Trem2 function culminates in a decrease in the number and activation of microglia in mouse models of AD or in mice treated with cuprizone to damage myelin (Cantoni et al., 2015; Ulrich et al., 2014; Wang et al., 2015). Trem2-deficient dendritic cells secrete more TNF-α, IL-6 and IL-12 compared with wild-type cells, particularly when activated with LPS suggesting there may be a shift towards cells expressing a pro-inflammatory and potentially more damaging phenotype in the absence of Trem2 (Hamerman et al., 2006; Turnbull et al., 2006). However, not all findings are consistent. A recent report demonstrated reduced AD pathology in an amyloid mouse crossed with a Trem2 knock-out mouse (Jay et al., 2015)

The main AD-associated TREM2 variant R47H has been shown to alter glycosylation and trafficking of the TREM2 protein between the golgi and ER resulting in fewer functional TREM2 receptors in the cell membrane and thus loss of TREM2 function (Park et al., 2015). The presence of this variant also reduces the cleavage of full-length TREM2 to a soluble extracellular fragment and, in both TREM2 risk variant carriers and in people with AD, less soluble TREM2 is present in CSF (Kleinberger et al., 2014) suggesting TREM2 dysfunction may be a common feature in AD and not just in those AD patients carrying a loss of function variant. Notwithstanding this, recent findings suggest that sTREM2 levels may be reduced or increased depending on the stage of AD and variant (Brendel et al., 2017).

TREM2 and other late onset AD susceptibility genes MS4A4A/4E/6A, CD33, HLA-DRB5/DRB1 and INPP5D are all part of a distinctive brain co-expression module which also contains the signaling partner for TREM2, TYROBP or DAP12 (Forabosco et al., 2013a; Hawrylycz et al., 2012; Zhang et al., 2013). This module appears to represent a biological network active in microglial cells with an innate immune function. It is significantly perturbed in AD brain (Forabosco et al., 2013b; Hawrylycz et al., 2012; Zhang et al., 2013) and remarkably contains fewer than 150 genes. This module shares identity with peripheral macrophages (Forabosco et al., 2013b) and many of the genes in the module are also altered in AD blood cells (Lunnon et al., 2012). Elevated expression of many of these genes, particularly TREM2 appears to be associated with the emergence of amyloid rather than Tau pathology in AD mouse models (Jiang et al., 2014; Matarin et al., 2015). Both TREM2 and Tyrobp have also been identified as major hubs in human APOE expressing mice following traumatic brain injury (Castranio et al., 2017).

Prominent voices in the field of AD research are proposing that to fully understand the etiology of AD we have to go beyond reductionist approaches and the amyloid cascade linearity and that there is a need for studies that address the complex cellular context of the disease which involves interactions between different cell types as the disease progresses across time and tissue (De Strooper and Karran, 2016). It is suggested that temporal resolution can be obtained from cohorts of mice at different stages of the disease. Previous studies using a systems level approach on TREM2 have lacked the aforementioned temporal resolution. Furthermore, the use of microarrays to measure gene expression could have meant that subtle effects are not detected. In our study, we tried to address these concerns by profiling Trem2 KO mice gene expression using RNA-Seq at two time points and tissues. We used brain cell markers to infer cell type specificity and detected gene co-expresion dysruption affecting a module with endothelial identity at an early stage that causes widespread disruption of other cell-type subnetworks, including a subnetwork with neuronal identity.

## 2 Methods

### 2.1 Design

Brain tissue samples were obtained from male Trem2 knockout (KO) and wild type (WT) control mice at two time points: 4 months and 8 months. These time points span the onset and late disease stages in well established AD mouse models (Matarin et al., 2015). Hippocampus and cortex were selected because they represent tissues affected in AD at early and late stages, respectively (Mastrangelo and Bowers, 2008; Matarin et al., 2015). RNA-Seq was used to profile the transcriptomes for each sample. Two technical replicates were obtained for each sample. Expression data analysed in this study are available at NCBI’s GEO through accession number GSE104381

Differential expression (DE) analysis allowed us to detect changes in expression between time points and tissues. Co-expression analysis was performed to detect higher level disturbances in gene expression networks. Enrichment analysis of the results allowed us to detect functions and pathways more altered in the absence of Trem2. Finally, the integration of cell type markers, enabled us to go further and not only detect time and tissue specific changes but uncover how the interactions between different cell types were affected and at which time point and tissue these changes were occurring. Fig. 1A displays an overview of the experimental and analytical workflow.

**Fig. 1.**
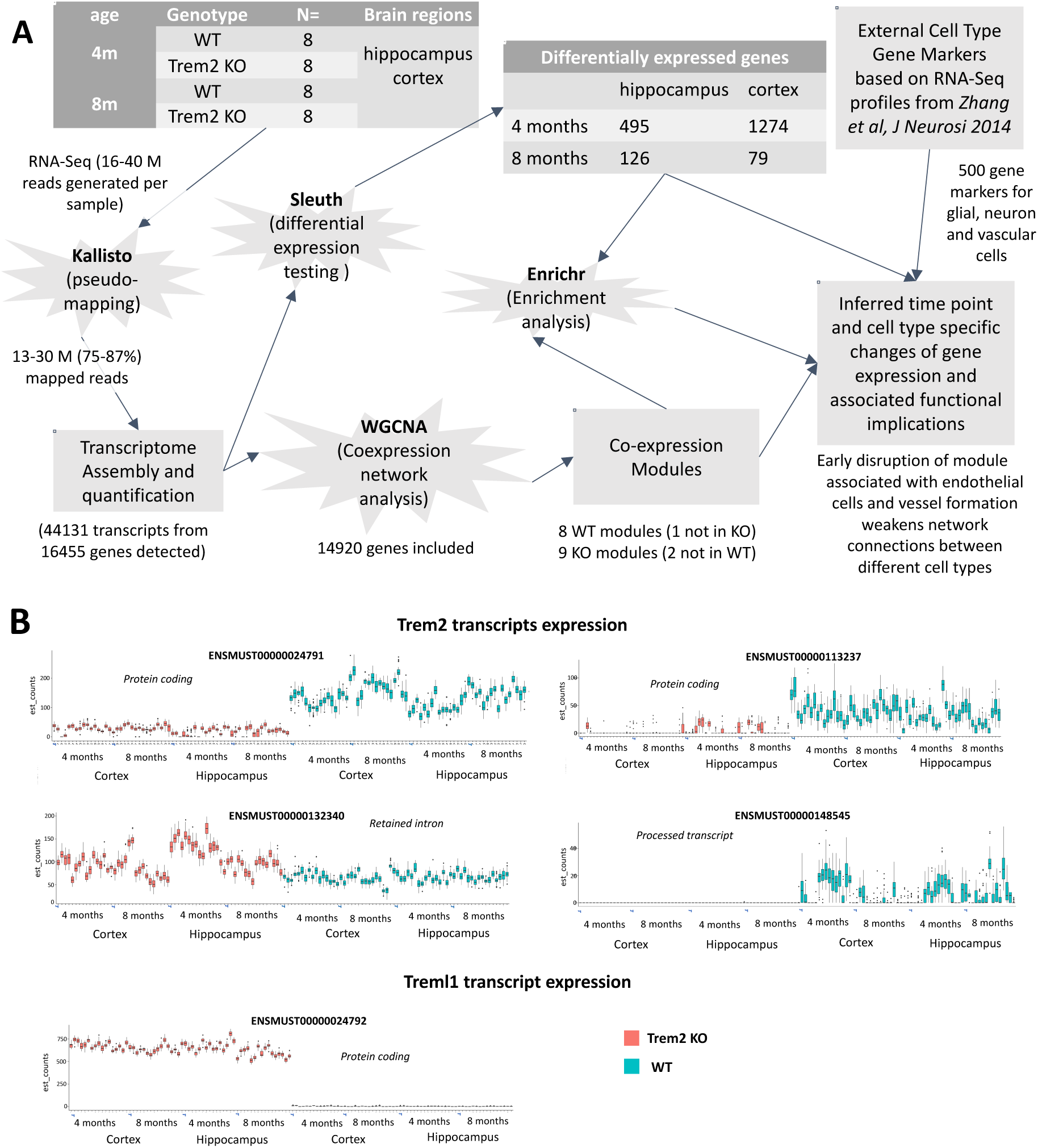
Experimental design, analysis work-flow and KO assessment. a) Experimental design and analysis workflow. Transcriptome profiling of 8 WT and 8 Trem2 knockout mice using RNA-Seq. Read alignment to quantify transcripts was done using Kallisto. Different expression analysis was done using Sleuth. Co-expression Network analysis was done using WGCNA. Enrichment analysis was done with Enrichr. In order to estimate cell type associations with co-expression modules gene markers were obtained from an external source\cite{Zhang2014}. All the results were contrasted to infer time and tissue specific changes caused by the absence of Trem2. b) Trem2 and Treml1 isoforms expression levels. Trem2 ensembl transcripts consist of two coding transcripts, ENSMUST00000024791 and ENSMUST00000113237, and two non-coding, ENSMUST00000132340 and ENSMUST00000148545. Treml1 only has one annotated coding transcript ENSMUST00000024792.

### 2.2 Generation of Trem2-/- mice

The Trem2-/- mouse model (Trem2tm1(KOMP)Vlcg), was generated by knocking a LacZ reporter cassette into the endogenous Trem2 locus in place of exons 2 and 3 and most of exon 4, resulting in a loss of Trem2 function as well as expression of the LacZ reporter under the control of the Trem2 promoter, as described previously (Jay et al., 2015). The mouse line was originally generated by the trans-NIH Knock-Out Mouse Project (KOMP). Frozen sperm were obtained from the UC Davis KOMP repository and a colony of mice was established at Taconic, Cambridge City USA. Mice were maintained on a B6 background and shipped to Eli Lilly and Co., Indianapolis, U.S.A. for subsequent tissue collection.

### 2.3 Sample description and RNA-Seq Library Construction

A total of 64 samples were used in the study. The hippocampus and cortex were dissected from the left hemisphere of each mouse brain, and snap frozen in liquid nitrogen. Tissues were homogenized in Qiazol reagent and total RNA isolated using the RNeasy Plus Universal Mini Kit according to the manufacturer’s protocol (Qiagen). RNA integrity (RIN) was measured for all samples using the Bioanalyser Agilent 2100 series using RNA nano chip. All sequencing libraries analyzed were generated from RNA samples measuring a RIN score ≥ 8.9. The Illumina TruSeq mRNA stranded protocol was used to obtain poly-A mRNA from all samples. 100ng of isolated mRNA was used to construct RNA-seq libraries. Libraries were quantified and normalized using the NEBNext^®^ mRNA Library prep kit for Illumina and sequenced as paired-end 76bp reads on the Illumina NextSeq 500 platform usiing technical replicates.

### 2.4 RNA-Seq Analysis

RNAseq reads were aligned to the mouse GRCm38 rel 79 reference transcriptome using Kallisto version 0.42.4 (Bray et al., 2016). The reference transcriptome was downloaded from the Kallisto project website (“http://bio.math.berkeley.edu/kallisto/transcriptomes/,” n.d.). A batch of quality control methods implemented in the R sleuth package were applied including PCA, MA plots, qq-plots and varplot(bootstrapping))(See Supplementary File 1). Quality control revealed consistent quality and no samples were removed. Differential gene expression analysis was performed on TPM normalized counts with the R sleuth package version 0.28.1 (H. Pimentel et al., 2016; H. J. Pimentel et al., 2016). Multiple biological replicates were used for all comparative analysis as well as a technical replicate for each of the samples. Significant genes, q-value ≤ 0.01, were used for gene ontology and pathway analysis. Gene ontology and pathway analysis was performed using the Enrichr database (Chen et al., 2013; Kuleshov et al., 2016; Newhouse, 2017) and Process Network Tool in *MetaCore*^*TM*^ (Thomson Reuters). Plots and graphics were generated using the sleuth (H. Pimentel et al., 2016; H. J. Pimentel et al., 2016) and ggplot version 2.2.1 (Wickham, 2009) R packages. R 3.3.0 version was used.

The number of reads per replicate ranged from 16 to 40 million reads (see Supplementary Table 1). The efficiency of the mapping varied from 75 to 87% with the number of reads mapped ranging from 13 million to 30 million. Each sample was profiled for 44,131 transcripts with 16,455 genes detected after quality control filtering. Principal component analysis revealed that samples clustered by age and tissue (see Supplementary File 1). Quality control revealed consistent quality and therefore all samples were used in subsequent analyses.

### 2.5 Gene network construction and module detection

For the co-expression analysis, we included all genes that were expressed in at least one time point, condition or tissue in the final set and with at least 10 reads in more than 90% of samples (Langfelder and Horvath, 2008). Using these criteria, Trem2 is retained for analysis, even though it is not expressed in the Trem2 KO mice. This left us with 14,920 genes (see Fig. 1A).

We used WGCNA version 1.51 to identify modules of co-expressed genes within gene expression networks (Langfelder and Horvath, 2008). To construct the network, biweight midcorrelations were calculated for all possible genes pairs. Values were entered in a matrix, and the data were transformed so that the matrix followed an approximate scale-free topology (see Supplementray File 2 for detailed information). A dynamic tree cut algorithm was used to detect network modules (Langfelder et al., 2008). We used signed networks to preserve the direction of correlations. We ran singular value decomposition on each module’s expression matrix and used the resulting module eigengene, which is equivalent to the first principal component (Langfelder and Horvath, 2007), to represent the overall expression profiles of the modules.

Cytoscape v3.5.1 was used to generate network maps of the most highly correlated genes and modules (Shannon et al., 2003).

### 2.6 Module matching between networks

Modules from the Trem2 KO network were matched to those of the WT network using a hypergeometric test to identify the WT module that has the most significant gene overlap and its colour was then assigned to the KO module.

### 2.7 Module preservation statistics

We applied both the module preservation Zsummary and the medianRank statistics to assess the module preservation from different expression data sets (Langfelder et al., 2011). Unlike the cross-tabulation test, Zsummary not only considers the overlap in module membership, but also the density and connectivity patterns of modules. In addition, for our study, network-based preservation statistics only require that module membership is identified in the original data sets, reducing the variation coming from various parameters setting to identify new modules in validation data sets. We converted the transcript-level measurements into gene-level measurements using the collapseRows R function (Miller et al., 2011). We adapted the method used to select the most representative probe to select the most representative transcript. The transcript isoform within a gene that had the highest average expression was used to represent that gene as it has been described that it leads to best between-study consistency. Overall, 14920 genes were retained in our preservation calculation.

### 2.8 Module membership statistics

Module membership measures correlations between each gene and each module eigenene, effectively measuring how strong is the connection of a gene with the module it has been assigned to and to the other modules too. We compared the correlation of the module membership values between the WT and KO networks using two approaches using the methods described by the WGCNA authors (Langfelder and Horvath, 2008). The first one, uses all the genes to assess overall conservation of the modules. The second one, only includes the genes assigned to the modules to assess "hub" conservation (i.e. are the genes more strongly correlated to a module in one of the networks also strongly correlated to the module in the other network).

### 2.9 Differential expression tests for modules

We used student t-tests to statistically compare expression differences between modules as suggested by the WGCNA authors (Langfelder and Horvath, 2008).

### 2.10 Integration of cell type markers

The top 500 most highly expressed genes for different brain cell types (neurons, endothelial cells, astrocytes, microglia, oligodendrocyte precursor cells, newly formed oligodendrocytes and myelinating oligodendrocytes) were selected from an RNA-Sequencing transcriptome database (Zhang et al., 2014). Of these, 411 astrocytes, 386 endothelial cells, 316 microglia, 407 myelinating oligodendrocytes, 394 newly formed oligodendrocytes, 390 oligodendrocyte precursors cells and 430 neurons marker genes could be mapped to our dataset and were used in subsequent analyses. The number of markers per cell type was divided by the total number of markers present for that given cell type to take into account for any bias in total number of cell markers available for different cell types. The obtained values were multiplied by 100 to avoid working with very small numbers.

### 2.11 Ethics statement

All animal procedures and experiments were performed in accordance with the Institutional Animal Care and Use Guidelines for Eli Lilly and Company. Protocol number 14-067.

## 3 Results

### 3.1 Analysis workflow and RNA-Seq assessment

In this study, we have applied an integrative approach for a systems level characterization of how the absence of Trem2 affects transcriptional expression (see the overview of the experimental and analytical workflow in Fig. 1A). This approach aims to make our results comparable with previous models of Alzheimer’s disease progression across tissue and time (Matarin et al., 2015). For this purpose, we used hippocampus and cortical regions to model progression across tissue and two time points, 4 and 8 months, to model progression with age. We compared the transcriptomes of WT and KO mice identifying differentially expressed genes and we used co-expression analysis to reveal changes in patterns of expression. To understand the functional implications of those changes, we integrated term enrichment analysis and brain cell type markers obtained from an external source (Zhang et al., 2014).

### 3.2 Trem2 knockout mouse assessment shows the Trem2 gene is effectively silenced

In order to assess the efficiency of the Trem2 knockout process, the levels of expression of the Ensembl (Ruffier et al., 2017) annotated Trem2 transcripts were explored (Fig. 1B). Both protein coding transcripts and a small processed transcript were silenced in the KO. A non-coding transcript containing a retained intron had higher expression in the KO, particularly at 4m in hippocampus. Treml1, the gene immediately downstream of Trem2, was silent in the WT but upregulated in the KO. The difference in Treml1 expression between WT and KO is the largest effect seen, even more so than that of Trem2, at all time points and tissues (Fig. 1B). It was recently described that this could be an artifact caused by the floxed neomycin selection cassette (Kang et al., 2017) employed in the generation of Trem2 knockout mice used for this is study. It is yet to determine if this artifact is functionally meaningful but we will interpret our results with caution.

### 3.3 The absence of Trem2 has a temporal effect on gene expression with a substantially bigger number of DE genes at 4m

Initially, we explored gene expression differences across 4 and 8 months aged mice (see Fig. 1A, Supplementary Files 5 and 6 and Supplementary Tables 2 to 5). A full list of differentially expressed genes is available in Supplementray Tables 2 to 5. We found that the number of DE genes between the WT and KO mice was 16 fold higher at 4 months cortex, where a total of 1274 genes were found to be differentially expressed, compared to 8 months cortex, where the expression of 79 genes was altered. The same trend was seen in the hippocampus, although fewer genes were differentially expressed at the early time-point - 495 at 4 months compared to 126 at 8 months.

The only substantial overlap in DE genes across time and tissue was found at 4 months, where the hippocampus and cortex shared 277 DE genes (see Supplementary File 6). This represents more than half of the DE genes in hippocampus but less than a quarter of the DE genes in cortex. Among these are genes previously associated with AD. For example Rbm3, which has been shown to mediate structural plasticity and have protective effects on cooling in neurodegeneration (Peretti et al., 2015), is downregulated in the KO. Nfya, which is also downregulated, has been described to have a role in angiogenesis (Xu et al., 2016). Treml2, which was upregulated in the cortex at 4 months and in both tissues at 8 months, has been reported to contain variants with a protective effect in Alzheimer’s disease (Benitez et al., 2014).

Besides the overlap at 4 months, there is a reduced number of genes that are differentially expressed in more than one time point and tissue (see Supplementary Tables 6 and 7). Only Trem2 (as expected), Treml1, Grp17 and Fam107a are differentially expressed at all time points and tissues. Interestingly, Grp17 has been previously associated with Nasu-Hakola disease (Satoh et al., 2017), a disease characterised by homozygous loss of function of Trem2 or its signalling partner Tyrobp (Paloneva et al., 2002), and has been successfully targeted with a marketed anti-asthmatic drug which reduces neuroinflammation and elevates hippocampal neurogenesis resulting in improvements to learning and memory in older animals (Marschallinger et al., 2015). Fam107a, also known as Drr1, has been described as is a stress-induced actin bundling factor that modulates synaptic efficacy and cognition (Schmidt et al., 2011) and has also been connected to brain cancer (Dudley et al., 2014; Le et al., 2010).

### 3.4 Gene ontology enrichment analysis of DE genes shows time and tissue specific enrichment of processes related to Alzheimer’s disease

Ontology analysis showed an enrichment for genes involved in response to unfolded proteins and ubiquitination related functions at 4 months in both tissues (Table 1; also see S8 Table). Both functions were driven mainly, but not exclusively, by the differential expression of genes of the DnaJ family (DnaJA2 and DnajB2) and heat shock proteins for example HspA8 and Hsph1. Enrichment for “response to heat” was significant in hippocampus and was driven by Camk2g and Camk2b, members of the calmodulin-dependent protein kinase subfamily involved in calcium signaling. A similar trend was seen in the cortex.

**Table 1:** Summary of ontology enrichment for the differentially expressed genes across time points and tissues. Adjusted p-values obtained using a Fisher exact test with Benjamini-Hochberg method for correction for multiple hypotheses using Enrichr (Chen et al., 2013)

| GO BP enriched term | Benjamini-Hochberg corrected p-value |  |  |  |
| --- | --- | --- | --- | --- |
|  | 4 months |  | 8 months |  |
|  | Hippocampus | Cortex | Hippocampus | Cortex |
| response to unfolded protein (GO:0006986) | 0.0003578 | 0.0001455 | 0.1678 | 0.1767 |
| ATF6-mediated unfolded protein response (GO:0036500) | 0.01047 | 0.002142 | NS | NS |
| regulation of cellular response to heat (GO:1900034) | 0.005745 | 0.0793 | 0.4287 | 0.3149 |
| regulation of protein ubiquitination (GO:0031396) | 0.01939 | 0.6671 | NS | 0.1239 |
| positive regulation of proteasomal ubiquitin-dependent protein catabolic process (GO:0032436) | 0.483 | 0.0007589 | NS | NS |
| SRP-dependent cotranslational protein targeting to membrane (GO:0006614) | 0.9078 | 1 | 4.71E-12 | 0.05501 |
| translation (GO:0006412) | NS | 1 | 2.27E-10 | 0.07171 |
| regulation of macroautophagy (GO:0016241) | NS | 0.6991 | 0.005104 | 0.3018 |
| organelle transport along microtubule (GO:0072384) | NS | NS | 0.01932 | NS |
| cellular response to vascular endothelial growth factor stimulus (GO:0035924) | 0.5601 | 0.4068 | NS | 0.06581 |

At 8 months both tissues showed enrichment for membrane targeting related functions, driven mainly by ribosomal proteins such as Rpl10, but there were also differences between them. In hippocampus, there was enrichment for “regulation of macroautophagy” driven by Uchl1, which has been previously implicated in Alzheimer’s disease, and also for “organelle transport along microtubule" driven by Cdc42 and CopG1. In cortex, enrichment for “cellular response to vascular endothelial growth factor stimulus” was driven by Flt1 (also know as VEGFR1) and Dll4, which have been both described as having a role in endothelial sprouting (Eilken et al., 2017; Pitulescu et al., 2017).

### 3.5 Trem2 WT and KO gene co-expression networks analysis identifies modules with cell type and/or functional identities

We generated two multi-tissue and multi-age networks, one for WT and one for the KO, and matched the latter to the former (see Methods and Fig. 2). The WT network consisted of 8 modules each containing between 127 and 6,477 gene members. The KO co-expression network comprised 9 modules ranging from 204 to 5,213 genes.

**Fig. 2.**
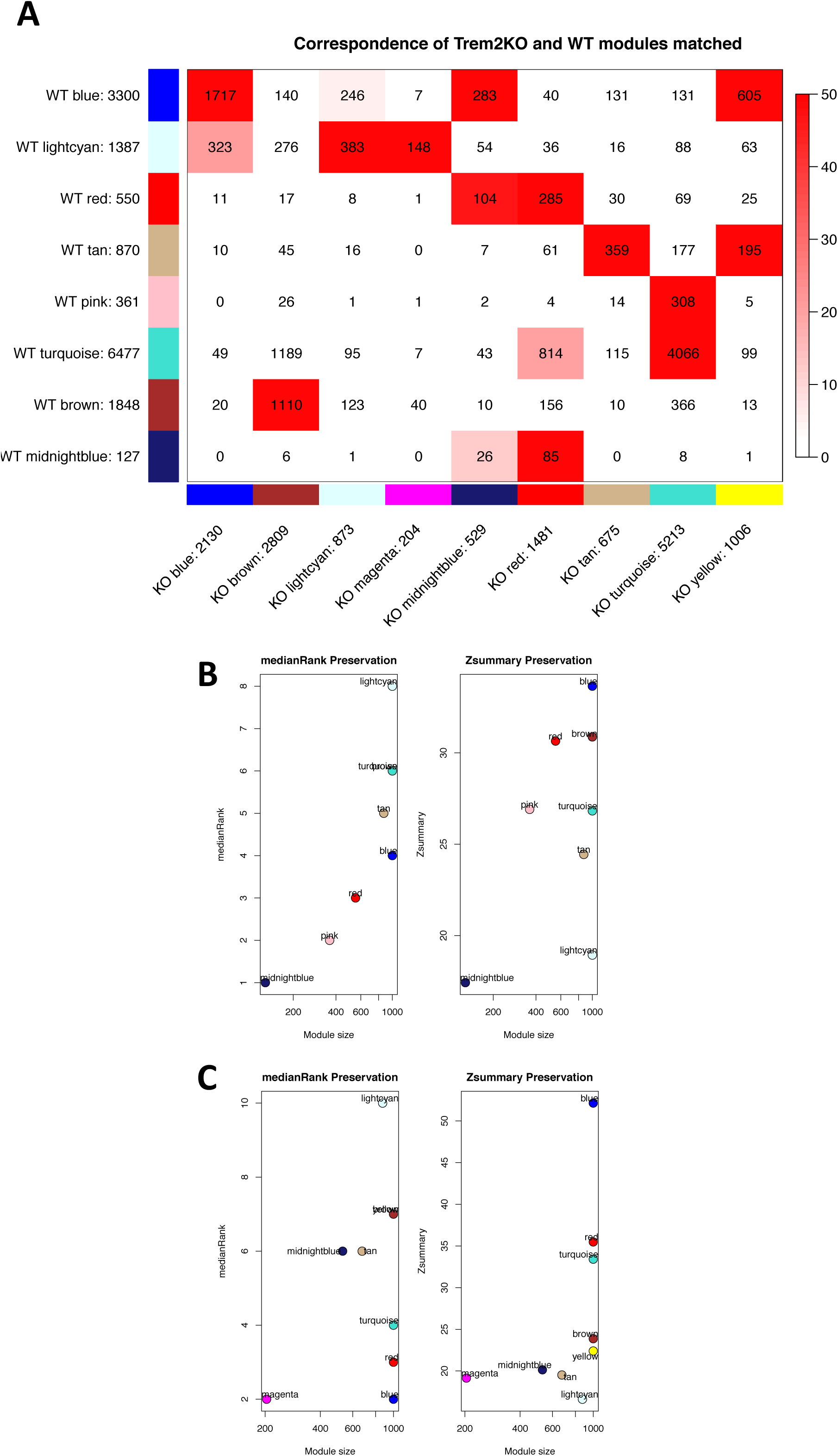
Module preservation between WT and Trem2 KO networks. a) Correspondence of WT set-specific modules and the Trem2 KO matched modules. Each row of the table corresponds to one KO set-specific module (labeled by color as well as text), and each column corresponds to one WT module. Numbers in the table indicate gene counts in the intersection of the corresponding modules. Coloring of the table encodes $-\log(p)$, with p being the hypergeometric test p-value for the overlap of the two modules. The stronger the red color, the more significant the overlap is. The table indicates that most WT set-specific modules have a KO counterpart. b) Preservation of WT modules in KO network. The left panel shows the composite statistic medianRank versus module size. The higher the medianRank the less preserved is the module relative to other modules. Since medianRank is based on the observed preservation statistics (as opposed to Z statistics or p-values) it is independent on module size. The right panel shows the composite statistic Zsummary. If Zsummary > 10 there is strong evidence that the module is preserved (Langfelder et al 2011). If Zsummary \textless 2, there is no evidence that the module preserved. Note that Zsummary shows a strong dependence on module size. c) Preservation of KO modules in WT network. The same as b) but in this case we assess the preservation of the modules obtained in the KO network to see how preserved they are compared to the ones obtained in the WT network.

Gene ontology and pathway analysis were used to assign biological significance to each module (see S9 Table). In the majority of cases, WT and KO modules were enriched for similar terms. In both WT and KO, the blue module was enriched for functions related to synaptic transmission and neurotransmitter secretion. Similarly, the brown module was enriched for mRNA transport and processing functions in both networks. The tan module was enriched for functions related to translation, protein targeting to membrane and mRNA catabolism. The turquoise module was strongly associated with ncRNA, mitochondrial related functions and house-keeping functions. The red module was associated with RNA splicing and cell cycle related functions. Conversely, the midnightblue module showed different functions in the WT and KO networks. In the WT, the module was associated with deubiquitination and, in the KO, most of its associated functions were related to synaptic transmission.

We used brain cell type markers (Zhang et al., 2014) as a proxy to test cell-type enrichment for each module (see methods and Figs 3 and 4). The WT and KO gene expression modules have similar cell type profiles (Supplementary File 3). The results of the enrichment for ontology terms, support those of the cell type enrichment analysis, for example, the blue module is enriched for neuron cell markers and for neuronal functions like synaptic transmission. The modules that had a clearer cell type identity were the mentioned blue module, with a neuronal identity, the lightcyan module with endothelial identity and the tan module with a microglial identity.

**Fig. 3.**
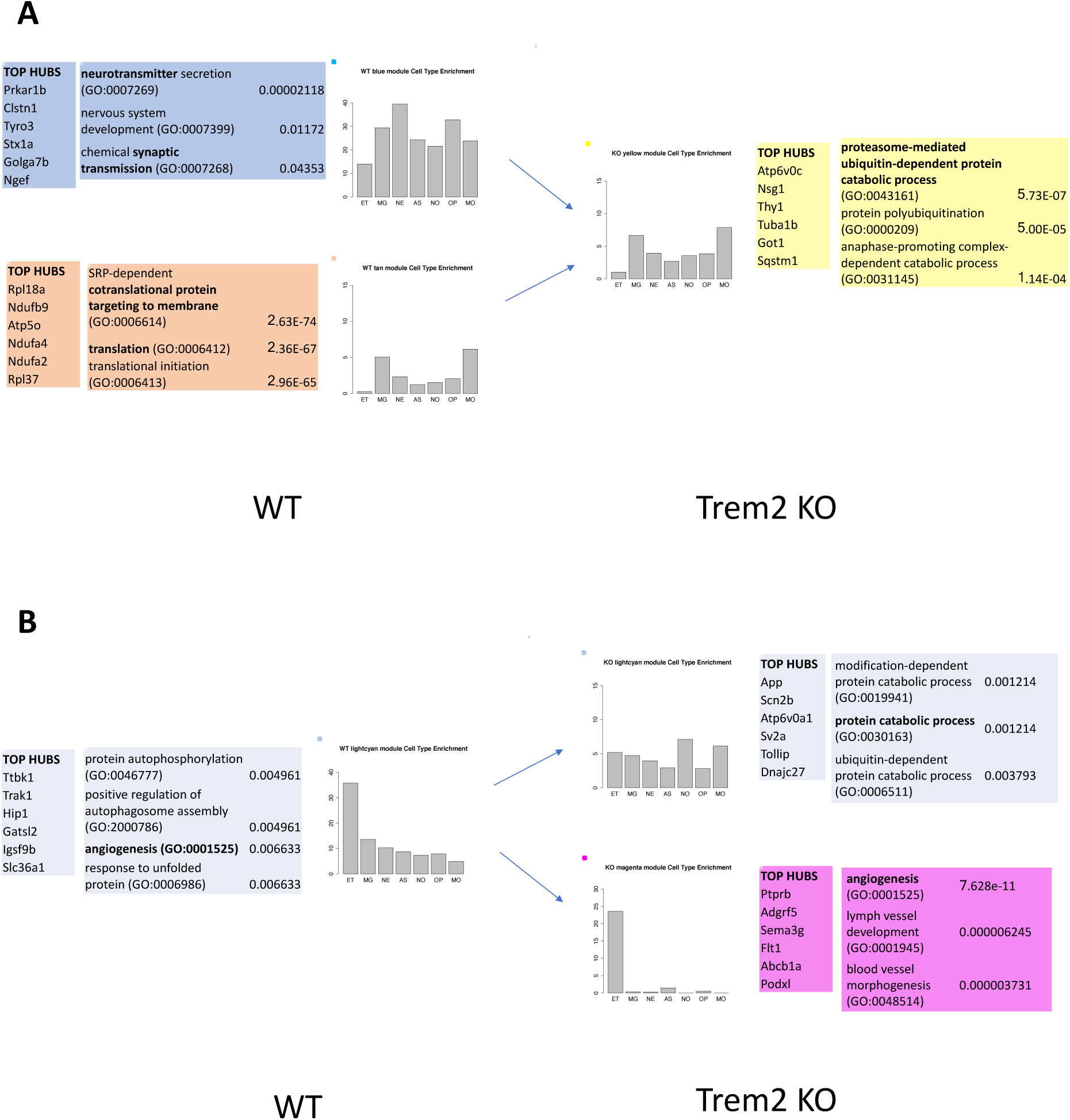
Summary of KO emerging modules top hubs and cell type and ontology enrichment. For each module considered, the list of the genes with the strongest correlation within its module, top hubs, is shown together with a list of ontology terms representative of the module function enrichment with its adjusted p-value (see Methods). The cell type identity of the network modules is inferred using gene markers enrichment. Each panel displays the proportion of markers for each cell type present in the module. The counts of the number of markers per cell type was divided by the number of markers present for that given cell type to take into account a possible bias (see methods for further details). The method allows us to approximate the proportion of each cell type that conform a module. For example, in the WT the blue module is predominantly associated with neuron markers and the lightcyan with endothelial cells. The cell types considered are astrocytes (AS), endothelial cells (ET), microglia (MG), myelinating oligodendrocytes (MO), newly formed oligodendrocytes (NO), oligodendrocyte precursors cells (OP) and neurons (NE). a) Emergence of KO yellow module by an increase in expression correlation of parts of the WT blue and tan modules. b) Emergence of KO magenta module by the disruption of the WT lightcyan module.

**Fig. 4.**
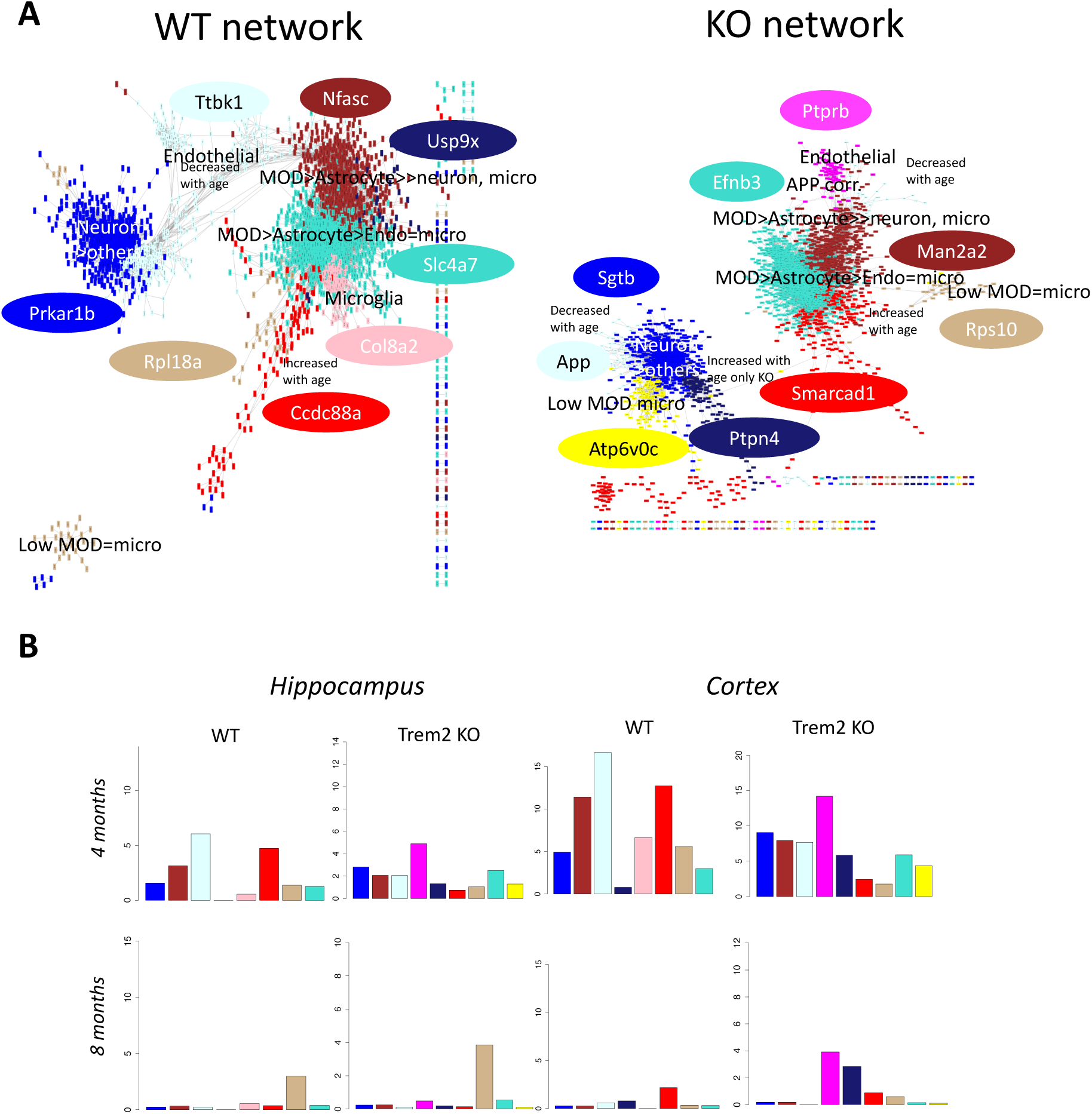
Disruption of the modules interconnectivity. a) Network view of the gene correlations higher than 0.5 in the WT (left panel) and the KO (right panel). Gene colours represent the module where they have been assigned. The lightcyan module, enriched for endothelial cells, is closely connected with the blue module, enriched for neurons, and in turn connects it to the glia enriched modules in the WT. In the KO, this connection is weakened when the majority of endothelia cell marker genes become members of a newly formed magenta module that is no longer strongly correlated with the blue module. The most correlated gene within each module, the top hub, is shown in an ellipse filled with the colour of the module it belongs to. The top hub of the lightcyan in the WT is Ttbk1, Tau Tubulin Kinase 1, which regulates phosphorylation of tau, and in the KO its place is taken by App, the gene that codes for the amyloid precursor, both proteins central to the Alzheimer’s pathology. The blue module expression association with age is decreased in the KO network but not in the WT (two tailed t-test with Welch modification, pval = 0.0036) and the midnightblue module expression is increase with age in the KO (two tailed t-test with Welch modification, pval = 0.017) but not in the WT. b) Differential expressed genes per module across time and tissue. Each panel contains the percentage of genes within a module that are differentially expressed (DE) respect the total number of genes in the module. The lightcyan module is the module with the higher proportion of DE genes at 4 months in both tissues in the WT. The magenta module, with can be considered a lightcyan spin-off since the majority of its genes have a counterpart in that module (see Fig 2A), is in turn the module with the higher proportion of DE genes in the KO. Differences in expression affect different modules in different tissues at 8 months, the tan module being the most affected in hippocampus while the changes in cortex are predominantly affecting the magenta module as it is also the case at 4 months.

#### 3.5.1 KO specific co-expression modules emerge notwithstanding broad conservation between WT and Trem2 KO co-expression networks

After identifying the modules in the two networks, we wanted to compare how well preserved they were. Given the cell type and functional assignments done previously we could potentially identify interesting functional or cell type perturbations between the two by looking at the differences between the network modules. Fig. 2A highlights the direct comparison of the WT and KO co-expression networks. It represents the number of genes for each module in each of the co-expression networks and the number of those genes that overlap with the modules in the other network. To formally assess the preservation of the WT modules in the KO network and vice versa, we calculated their Zsummary preservation scores and medianRank preservation values (Fig. 2B and Fig. 2C) (Langfelder et al., 2011) (see Methods section).

Most WT modules have a KO counterpart, with the exception of the WT pink module. Similarly, all the KO co-expression modules had a WT counterpart, except the KO magenta and yellow modules.

The WT lightcyan module was ranked as the least preserved module. The WT tan module had one of the lowest Zsummary scores and rankings, as does its KO counterpart. The tan module was of particular interest because it contained Trem2 and its signaling partner Tyrobp and, consistenly with Trem2 being expressed in microglia, was assigned a microglial identity using cell type markers.

The KO magenta, midnightblue, yellow and brown had relatively lower Zsummary values but we could not consider any module as not "preserved" since they all had Zsummary scores higher than 10 (Horvath, 2011).

Zsummary scores were influenced by module size which can explain the apparent contradiction of the midnightblue module being at the same time the best ranked for preservation and the one with the lower Zsummary score. This module is markedly different between the WT and KO, partially because the colour assignment of the KO midnightblue module is based on 26 overlapping genes (only 5% of the 529 genes in the module) with the WT midnightblue module.

To further study the module preservation between the two networks we used module membership (MM) values. The modules with the lowest correlations in module membership between the WT and KO were the lightcyan and tan modules (see Supplementary Files 7 and 8), the same ones that had overall lowest preservation scores. MM effectively measures how strong is the connection of a gene with the module it has been assigned to, calculating the correlation between the gene and the module eigenene. The genes with the highest MM are the most highly connected genes within the module and are referred to as "hubs". This correlation is calculated not only for the module it has been assigned to but for the other modules too so it is useful to compare different networks where the genes may have been assigned to different modules (more details in the methods section). Despite broad conservation, there were modules that could not be matched between the two networks. The WT pink module had no KO counterpart, but it had a relatively high Zsummary score compared to the rest of the modules(see Fig. 2B) and was one of the modules with the highest module membership correlation with the KO network (cor 0.89, p-value<0.01; see Supplementary File 8). Interestingly, it was enriched for extracellular matrix organization and exosome-related functions and both endothelial and microglia cell type markers. Furthermore, the pink module top hub gene Col8a2, encodes a protein whose absence has been described to produce progressive alterations in endothelial cell morphology and cell loss (Jun et al., 2012). Npc2, one of the genes causing frontotemporal dementia when mutated (Zech et al., 2013), is also part of this module.

In the KO network, the magenta and yellow modules were not matched to any WT modules and had low Zsummary scores. 72% of magenta module genes overlap with the lightcyan-endothelial module, on eof the least preserved modules. Nearly 20% of the genes of the yellow module ovelap with the tan module, the other least preserved module, and 60% of them overlap the blue module with neuronal identity. We considered that this deserved further inspection and we will look at them in more detail in the following sections.

#### 3.5.2 The KO yellow module is enriched for proteosome catabolic processes, microglia cell markers and genes previously implicated in different dementias

The KO yellow module contained genes overlapping mostly with the WT blue and WT tan modules (see Fig. 2A). We looked at the different gene ontology and cell type enrichment profiles and top hub genes of these modules to further understand the yellow module emergence in the KO (see Fig. 3A).

The WT blue module was enriched for neuron cell markers and terms associated with neuronal cell functions, such as neurotransmitter secretion and synaptic transmission. Its top hub gene Prkar1b has been previously linked with neurodegeneration (Wong et al., 2014) and the second one, Clstn1, has been shown to be associated with pathogenic mechanisms of AD (Vagnoni et al., 2012).

The WT tan module was enriched for microglia cell type markers and for translation related ontology terms and its top hub gene was the ribosomal protein, Rpl18a.

The KO yellow module that emerged as parts of the WT blue and WT tan module had a cell marker profile very similar to the WT tan module (microglia) but was also enriched for proteasome, ubiquitin and catabolism related processes.

Interestingly, the KO yellow module top hub gene was Atp6v0c. This gene encodes a component of vacuolar ATPase (V-ATPase), a multisubunit enzyme that is necessary for intracellular processes receptor-mediated endocytosis, and synaptic vesicle proton gradient generation (Lee et al., 2010). Atp6v0c has been shown to alter autophagy-lysosome pathway function and metabolism of proteins that accumulate in neurodegenerative disease (Mangieri et al., 2014) and has been described as a promising target for therapeutic development (Higashida et al., 2017). Other genes previously associated with different forms of dementia form part of this module. Mapt, the gene that encodes the tau protein which has been associated with numerous types of neurodegenerative disorders (Spillantini and Goedert, 2013), becomes strongly associated with the module. Pink1, a gene that causes Parkinson’s disease when mutated, is also strongly associated, as it is Sqtsm1, a gene involved in sporadic amyotrophic lateral sclerosis pathology (Fecto, 2011).

Altogether, the characteristics of the yellow module relate to known mechanisms involved in the pathology of different types of dementias.

#### 3.5.3 The KO magenta module is enriched for endothelial cell markers and functions

The KO magenta module emerged from the disruption of the WT lightcyan module as can be seen from the module overlap in Fig. 2A. Over 72%, 148 out of 204, of the genes in the KO magenta module overlapped with those of the WT lightcyan module. As we did before for the yellow module, we looked at the different enrichment profiles and top hub genes of these modules to further understand the module emergence in the KO (see Fig. 3B).

The WT lightcyan module was enriched for endothelial cell markers and terms associated with endothelial cell functions, like angiogenesis, and also for other functions related to AD such as phagocytosis and response to unfolded protein. Its top hub gene Ttbk1(tau tubulin kinase 1), whose encoded protein is a neuron-specific serine/threonine and tyrosine kinase that regulates phosphorylation of tau, has been implicated in both the pathology of amyotrophic lateral sclerosis (Liachko et al., 2014) and AD (Vázquez-Higuera et al., 2011).

When the lightcyan module broke up its KO counterpart became enriched for ubiquitin and catabolism related processes. The WT hub gene Ttbk1, lost its hub status in the KO module, where the top hub became App, the amyloid precursor gene. It also worth mentioning that in the KO one of the top hub genes, the third one (see Fig. 3B) is Atp6v0a1, which is one of the components of the vacuolar ATPase together with Atp6v0c, the top hub of the yellow module.

The module was not enriched for endothelial cell type markers anymore but shifted towards an oligodendrocyte identity. This was caused by the vast majority of endothelial cell markers moving to the new magenta module, which was significantly enriched for genes with blood vessel development function. The magenta module top hub, Ptprb, is also reportedly involved in the establishment of endothelial cell polarity and lumen formation (Hayashi et al., 2013). Interestingly, Flt1 (also known as vascular endothelial growth factor receptor 1) was another top hub in the magenta module. Flt1 has been described to have a role in motor neuron degeneration (Poesen et al., 2008). Most importantly, the disruption of the lightcyan module in the KO network suggested that endothelial cells become dyscoordinated in the absence of Trem2.

#### 3.5.4 Dyscoordination of the endothelial module weakens neuron module and glia module coexpression

To help us visualise better the differences between the wild type and knockout networks we plotted the co-expression networks with genes coloured by their module assignment including only correlations stronger than 0.5 (see Fig. 4A). In this way, we could see how the disruption of the endothelial module in the KO network weakens the connection of the whole network in general and in particular the connections between the neuronal subnetwork and the modules with glial identity.

In the WT, the lightcyan module (enriched for endothelial cell functions and markers) was connected on one end with the blue module (enriched for neuron functions and cell markers) and, on the other end, the lighcyan-endothelial module connected with other modules enriched for glia markers and functions. In particular the WT endothelial subnetwork was connected with the brown module, which is mainly enriched for myelinating oligodendrocytes and astrocytes. In the KO, these connections were weakened by the endothelial module disruption. The KO lightcyan module counterpart remained strongly connected to the neuron module and App became its the top hub gene. The KO specific magenta module, which has the biggest proportion of endothelial gene markers lost to the lightcyan module in the KO, lost the strong connection to the neuron module but remained correlated with the brown module.

Interestingly, the new KO yellow module, enriched for proteosome functions and genes previously associated with different types of dementias, was also strongly correlated to the neuron module.

We can also see in Fig. 4A how the midnightblue module is markedly different between the WT and KO and is strongly associated with the blue neuronal module in the KO. The KO midnightblue module was enriched for neuron markers but also for the gamma-aminobuyiric acid (GABA) signaling pathway (see S9 Table). While the blue module expression decreases with age in the KO (t-test with Welch correction, p = 0.004), the expression of the midnightblue increases (t-test with Welch correction, p = 0.02)(see Supplementary File 4). Interestingly, GABA signaling has been described to be increased in 5xFAD Alzheimer’s mouse models and its reduction has been described to rescue the impairment of long-term potentiation (LTP) and memory deficit (Wu et al., 2014).

Mapping the differentially expressed genes to the modules obtained in the previous section we were able to visualise which modules are affected at different ages in different tissues (see Fig. 4B). At 4 months, most of the changes in expression in both hippocampus and cortex affected the lightcyan module in the WT or the magenta module, product of the WT lightcyan disruption, if we look at the KO network. This points to the endothelial module disruption being an early event. The lightcyan module is enriched for response to unfolded protein GO terms in the WT and for ubiquitination in the KO. This is consistent with what we found in the enrichment analysis for genes being differentially expressed at 4 months (described in Table 1). At 4 months, the rest of the modules were broadly affected to a similar extent in both tissues. Notable exceptions are the WT specific pink module affected specifically in the cortex at 4 months and the yellow KO module that shows larger differences at 4 month in cortex, although not as dramatically as the pink module.

At 8 months, there was a very different picture with changes in expression affecting different modules in different tissues. The tan module is the more affected in the hippocampus. Again, consistent with the enrichment analysis performed previously for the differentially expressed genes, we find enrichment for macroautophagy in both the tan module and list of differential expressed genes at 8m in hippocampus. The magenta endothelial module is the most affected in the cortex. This again agrees with the enrichment for “cellular response to vascular endothelial growth factor stimulus” specifically in the list of differentially expressed genes at 8 months cortex.

Overall, we found that the absence of Trem2 in microglia produces a ripple effect that provokes the disruption of a subnetwork of genes enriched for endothelial cell gene markers and functions at an early time point, 4 months of age, with the effect being stronger in the cortex at this early time point. At a later stage, 8 months of age, cortex and hippocampus are affected differently with the subnetwork of genes with microglial identity being more perturbed in the hippocampus. This shows that the absence of Trem2 has time and tissue specific effects that evolve with age.

## 4 Discussion

Many risk genes contributing to dementia appear to have pleiotropic effects Rare loss of function variants in the TREM2 gene can cause Nasu-Hakola disease when both alleles are affected (Paloneva et al., 2000) or increase the risk to develop Alzheimer’s disease, Parkinson's disease or Frontotemporal dementia in heterozygous carriers (R. Guerreiro et al., 2013; Jonsson et al., 2013). Further study of the complex molecular consequences of the absence of TREM2 could have a big impact on our understanding of AD pathogenesis from a systems level view (De Strooper and Karran, 2016). We generated RNA-Seq data to profile hippocampus and cortex at 4 and 8 months of age to explore the temporal and spatial consequences of TREM2 absence. Differentially expressed genes were enriched for functions related to Alzheimer’s disease, and other forms of dementia. Furthermore, WGCNA analysis coupled with enrichment for cell type markers highlighted the disruption of a co-expression module with a strong endothelial identity. The absence of Trem2 in microglia appears to generate a ripple effect that causes the disruption of an alignment betwen endothelial cells and microglia affecting the coordination of neurons and glia. Interestingly, this suggests a new role for Trem2 beyond its known functions as a microglial receptor and signalling hub, linking the immune response with vascular consequences as potential underlying causes of Alzheimer’s disease.

Our study involved expression profiling of 8 Trem2 knockout and 8 wild type mice using RNA-Seq. We sampled hippocampus and cortex at 4 months and 8 months of age. This allowed us to model the disease progression across tissue and time and enables results to be compared with previous expression profiling of transgenic mice that develop amyloid or tau pathology (Matarin et al., 2015). Surprisingly, the level of expression of Trem2 in the wild type mice increases from 4 months to 8 months in both tissues, at least for the most highly expressed isoform. The differences may be attributed to the use of RNA-Seq which let us consider each isoform independently. In fact, of the two protein coding isoforms included in Ensembl we only detect significant differential expression between 4 months and 8 months for Trem2-201. As previously noted, microglia exclusively express TREM2 in the brain (Ulrich et al., 2017) and the increased levels of expression may reflect an increase in microglia density with age (Conde and Streit, 2006). Other differential effects between 4 and 8 months were also found. Nearly 1,300 genes had significant differential expression at 4 months which was reduced to only 79 genes by 8 months. Interestingly, the number of DE genes was greater in hippocampus than in cortex at 8 months suggesting a shift in dominance in TREM2 activity between brain regions. These alterations follow a regional distribution that is consistent with that of Nasu-Hakola, whose pathology affects centre in the cortical region earlier in life than AD which has a much later onset and centres at least initially in the hippocampus (Ridha et al., 2004)

Our WGCNA coexpression analysis coupled with cell brain cell type markers (Zhang et al., 2014) used as a proxy of cell-type enrichment helped us to identify modules with both functional and cell type identities. We found a module, blue, enriched for synaptic transmission ontology terms had the highest enrichment for neuronal markers. We also identified a module that contained Trem2 and Tyrobp and was mostly enriched for microglia and myelinating oligodendrocytes. The module containing Trem2 and Tyrobp in our study is larger and has only modest gene overlap compared to what was found in previous brain coexpression network analyses (Forabosco et al., 2013a; Zhang et al., 2013). Differences are to be expected. Previous studies involved the analysis of postmortem human control and/or AD brain tissue, and in both cases expression was profiled using microarray data. Nevertheless, some of the key genes such as Fcer1g and Ly86 identified in the AD immune module (Zhang et al., 2013) and Cxcr1 are present in the Tyrobp module in our study or, as is the case of Cxcr1, in the phagocytic yellow module that is specific to the knockout network.

This emerging KO yellow module which draws its genes from the module with neuronal identity and the module containing Trem2 and Tyrobp tan was enriched in catabolic, proteasome and degradation associated functions. Interestingly, its top hub is Atp6v0c, a gene that encodes a component of vacuolar ATPase (V-ATPase), a multisubunit enzyme that is necessary for intracellular processes receptor-mediated endocytosis and synaptic vesicle proton gradient generation (Lee et al., 2010), and has been described to alter autophagy-lysosome pathway function and metabolism of proteins that accumulate in neurodegenerative disease (Mangieri et al., 2014) and has been proposed as a possible target gene for therapy (Higashida et al., 2017). Furthermore, a number of genes that have previously associated with dementia form part of this module. Mapt, the gene encoding for the tau protein associated with different types of neurodegenerative disorders (Spillantini and Goedert, 2013), Pink1, a gene that can cause Parkinson’s disease when mutated and Sqtsm1, a gene involved in sporadic amyotrophic lateral sclerosis pathology (Fecto, 2011), are all part of this module suggesting that all these neurodegenerative pathologies may share and underlying molecular mechanism in which Trem2 plays a central role.

The WT lightcyan module (enriched for phosphorylation regulation, angiogenesis and protein processes and junction-related cellular compartments) was disrupted in the KO, showing the lowest preservation and breaking up to form the magenta module as a consequence. The KO specific magenta module was enriched in angiogenesis, similarly to the WT lightcyan module with whom it shares most of its genes, but it not was enriched for phosphorylation regulation. The top hub gene in the WT lightcyan module was Ttbk1 (tau tubulin kinase 1) but in the disrupted KO module App, the amyloid precursor protein, becomes the top hub status, while the top hub in the magenta module is Ptprb, which plays an important role in blood vessel remodelling and angiogenesis. This disruption suggests that Trem2 provides a link between endothelial cells and neurons that appears essential to the survival of neurons. When the coordination between the two falters App drives a subnetwork of genes involved in catabolic processes that is closely associated with the neuronal subnetwork. This finding also points to a role of Trem2 in the mechanisms behind the increased risk of dementia caused by vascular damage (Mantzavinos and Alexiou, 2017) and could also help explain the brain blood vessel damage described in cases of Nasu-Hakola disease (Kalimo et al., 1994)

The changes affecting the module with endothelial identity described previously occurred mainly at 4 months in both tissues, but the effect was strongest in cortex. These findings are consistent with human studies on Alzheimer’s disease (AD) where early changes are described in the prefrontal cortex region, for example changes in monoamine oxidase A and B (Kennedy et al., 2003) and these correlate with Braak stage (Braak and Braak, 1997). At 8 months, the changes are different from 4 months. A more phagocytosis-dominated module is altered in the hippocampus perhaps coinciding with the emergence of AD pathologies such as plaque deposition. Somewhat surprisingly, a number of complement genes are differentially expressed at 4 months in this module, C1qb in hippocampus and C1qa in cortex, but not at 8 months, where they lose their previous association with the module having neuron identity. The complement system becomes activated in response to the presence of amyloid and is therefore a pathological hallmark of AD (Lui et al., 2016).

To conclude, we have found that the timing and magnitude of the effect of Trem2 absence is different for different brain regions. Overall there were more changes in the cortex in younger mice capturing what is observed in Nasu-Hakola disease, which is characterised by frontotemporal dementia and blood vessel dysfunction (Kondo et al., 2002), while the hippocampus in older mice had more changes mirroring (R. Guerreiro et al., 2013; Jonsson et al., 2013)the vulnerability of the hippocampus in later onset Alzheimer’s disease (Mastrangelo and Bowers, 2008; Matarin et al., 2015). WGCNA analysis and brain cell type marker integration revealed links between several molecular processes previously described to play a part in AD pathology. In the absence of Trem2 a central role for amyloid processing emerged, as did changes in phagocytosis and altered vascularity. This reflects a more complex interplay between an absence of TREM2 and it's effects in different tissue and time and underlines the importance of considering the impact of disease and risk genes on the whole brain network when exploring cause, effect and developing treatments. It will be important to see if these findings can be replicated in people carrying the R47H risk variant associated with increased risk for Alzheimer’s disease (R. Guerreiro et al., 2013; Jonsson et al., 2013).

## Acknowledgements

AKH, GC, RJBD and SJN are supported by the the Institute of Psychiatry, Psychology and Neuroscience, King’s College London, London, UK. The project is co-funded by the Eli Lilly and Company Lilly Research Award Program (LRAP) which also supports GC. KM, NL, DAC and MJON are employees of Eli Lilly and Company Ltd, UK. JR, HW are employees of Eli Lilly and Company (Indianapolis, USA).

RJBD and SJN are also part of NIHR Biomedical Research Centre at South London and Maudsley NHS Foundation Trust and King’s College London, UK and Farr Institute of Health Informatics Research, UCL Institute of Health Informatics, University College London, London, United Kingdom. This project has also received funding from the Innovative Medicines Initiative 2 Joint Undertaking under grant agreement No 115976. This Joint Undertaking receives support from the European Union’s Horizon 2020 research and innovation programme and EFPIA. RJBD, SJN are also supported by the National Institute for Health Research (NIHR) University College London Hospitals Biomedical Research Centre, and by awards establishing the Farr Institute of Health Informatics Research at UCL Partners, from the Medical Research Council, Arthritis Research UK, British Heart Foundation, Cancer Research UK, Chief Scientist Office, Economic and Social Research Council, Engineering and Physical Sciences Research Council, National Institute for Health Research, National Institute for Social Care and Health Research, and Wellcome Trust (grant MR/K006584/1).

## Appendix A. Supplementary data

Supplementary File 1. QC plots for mapping data. QC plots for kallisto read mappings. Includes PCA plots showing clustering by and tissue and MAplots, qqplots, variance and volcano plots for the WT vs KO comparisons for different age and tissue groupings (4 month cortex, 4m hippocampus, 8 month cortex and 8 month hippocampus).

Supplementary File 2. Samples clustering and scale free topology fit for WT and KO networks. Plots showing clustering of the samples by tissue, age and individual mouse. Plots showing how well a scale free topology is fitted for WT and KO generated networks.

Supplementary File 3. Cell type enrichment for each module for WT and KO networks. Enrichment for cell type gene markers for each of the individual modules identified in the WT and KO networks.

Supplementary File 4. Age related module differential expression for WT and KO networks. Box plots comparing the levels of expression of the genes contained within each module at different ages to identify how module expression changes with age and t-test results to assess the differential expression between the two time points. Includes WT and

KO matched modules to identify differences between WT and KO networks.

Supplementary File 5. Venn diagram of differential expression gene overlap. Venn diagram describing the gene overlap between the different differential expression calls between different tissues and time points

Supplementary File 6. Interactive Venn diagram of differential expression gene overlap. Venn diagram of differential expression gene overlap data file that can be visualized by uploading it to http://www.interactivenn.net. It gives details about the overlapping genes that can be obtained by clicking on every overlapping section of the diagram.

Supplementary File 7. Module membership correlation between modules in the WT network and the colour matched modules in the knockout network. Includes all the matched genes used to generate the networks to assess global module conservation.

Supplementary File 8. Module membership correlation between modules in the WT network and the colour matched modules in the knockout network. Includes only the matched genes used to generate the networks that are assigned to each module to assess hub conservation.

Supplementary Table 1. Summary of mapping statistics per sample. Description of the mapping statistics of the mappings done with kallisto including number of reads mapped, number of reads processed, fraction of reads mapped and rounds of bootstrapping for each sample. Includes covariates for easier inspection of the results.

Supplementary Table 2. Complete list of DE gene calls between WT and KO for 4 month cortex.

Supplementary Table 3. Complete list of DE gene calls between WT and KO for 4 month hippocampus.

Supplementary Table 4. Complete list of DE gene calls between WT and KO for 8 month cortex.

Supplementary Table 5. Complete list of DE gene calls between WT

Supplementary Table 6. WT network module membership summary. For each of the genes included in the analysis, description of the WT modules module membership statistics and differential expression statistics (b and p-values) for each of the different WT vs KO comparisons.

Supplementary Table 7. KO network module membership summary. For each of the genes included in the analysis, description of the KO modules module membership statistics and differential expression statistics (b and p-values) for each of the different WT vs KO comparisons.

Supplementary Table 8. Gene ontology (GO) biological process (BP) enrichment for the different WT vs KO comparisons. Details including statistics of the Gene Ontolology enrichment analysis for each of the lists of genes obtained in the different comparisons done in the differential expression analysis.

Supplementary Table 9. Gene ontology (GO) biological process (BP) enrichment for the WT and KO modules. Details including statistics of the Gene Ontolology enrichment analysis for each of the lists of the genes contained in each of the modules obtained for the WT and KO networks in the WGCNA co-expression analysis.

